# Synaptotagmin-1 C2B domains cooperatively stabilize the fusion stalk via a master-servant mechanism

**DOI:** 10.1101/2021.11.29.470409

**Authors:** Ary Lautaro Di Bartolo, Diego Masone

**Author notes:** Electronic Supplementary Information (ESI) available: supplementary material contains umbrella sampling convergence analysis; time-averaged densities for stalk formation; molecular dynamics snapshots in the *μ*s scale; lipid species density profiles; membrane curvature measurements; inter-membrane lipid count over unbiased simulations and example input parameters for PLUMED. See DOI: 00.0000/00000000.

## Abstract

Synaptotagmin-1 is a low-affinity Ca^2+^ sensor that triggers synchronous vesicle fusion. It contains two similar C2 domains (C2A and C2B) that cooperate in membrane binding, being the C2B domain the main responsible for the membrane fusion process due to its polybasic patch KRLKKKKTTIKK (321-332). In this work, a master-servant mechanism between two identical C2B domains is shown to control the formation of the fusion stalk. Two regions in C2B are essential for the process, the well-known polybasic patch and a recently described pair of arginines (398,399). The master domain shows strong PIP_2_ interactions with its polybasic patch and its pair of arginines. At the same time, the servant analogously cooperates with the master to reduce the total work to form the fusion stalk. The strategic mutation (T328E,T329E) in both master and servant domains disrupts the cooperative mechanism, drastically increasing the free energy needed to induce the fusion stalk, however with negligible effects on the master domain interactions with PIP_2_. These data point to a difference in the behavior of the servant domain, which is unable to sustain its PIP_2_ interactions neither through its polybasic patch nor through its pair of arginines, in the end losing its ability to assist the master in the formation of the fusion stalk.

## 1 Introduction

Exocytosis is an important process used by eukaryotic cells to release biological compounds and transport lipids and proteins through the plasma membrane. Specialized secretory cells experience regulated exocytosis as a response to physiological signals ^1–3^. Mainly, sperm exocytosis (or acrosome reaction) is a regulated secretion needed to fertilize the egg that requires large membrane remodeling, membrane bending and fusion ^2,4–7^. While this collective process develops, multiple fusion pores spontaneously form between the acrosomal and plasma membranes, connecting the acrosome lumen to the extracellular milieu. Consequently, the fusion pore works as a remarkable mechanism to connect intracellular organelles and release vesicle contents during exocytosis. Figure 1 schematically shows membrane remodeling and fusion stalk formation between organelles, which is the first step in the formation of a fusion pore.

**Fig. 1.**
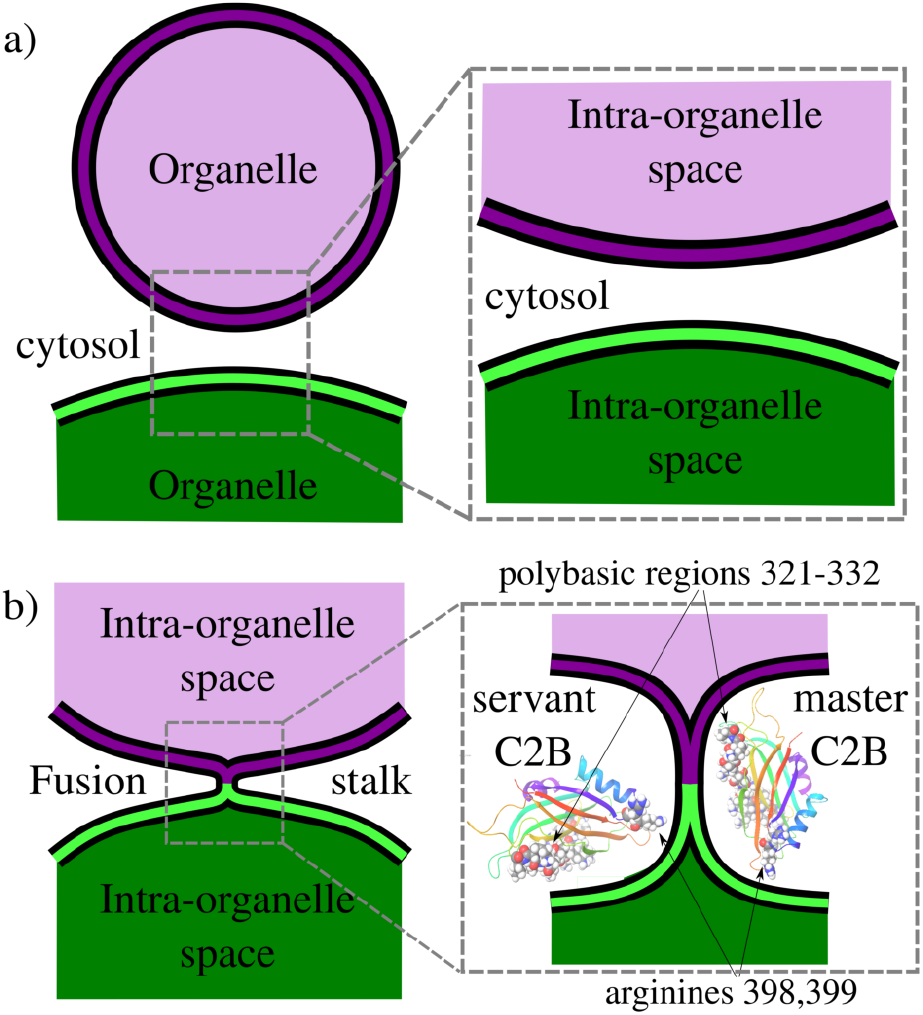
Schematics of the fusion stalk. a) Organelles about to fuse in the cytosol. b) Formation of the fusion stalk while interacting with Syt1-C2B domains. Arginines R398,R399 and the polybasic region 321-332 are highlighted in vdW representations.

Synaptotagmin-1 (Syt1) is a vesicle-anchored protein known as a phospholipid binding machine ^8^. Syt1 has been related to synaptic vesicle fusion ^9^, fusion pore opening ^10^, stabilization ^5,11^ and expansion ^12,13^. Syt1 contains two C2 domains (C2A and C2B) with Ca^2+^ binding loops, the latter (C2B) with a polybasic region KRLKKKKTTIKK (positions 321-332) that easily binds to negatively charged membrane patches, such as clusters of phosphatidylinositol-4,5-bisphosphate lipids (PI(4,5)P_2_, or simply PIP2), independently of Ca^2+ 14–18^. Therefore, the Syt1-C2B domain has been identified as the main energetic driver during membrane fusion and evoked neurotransmitter release ^19^. Importantly, Cafiso and collaborators ^13^ showed that two different regions of the C2B domain make unique contributions to the fusion process, namely, the polybasic region and a pair of arginines (R398,R399). However, the molecular mechanism by which Syt1 drives membrane fusion is yet not completely understood ^13,20^.

In the present work, we use enhanced molecular dynamics with an *ad-hoc* collective variable to induce a fusion stalk between lipid bilayers. We demonstrate that Syt1-C2B domains cooperatively facilitate the formation of the fusion stalk, significantly reducing its total thermodynamic work. We observe a master-servant mechanism between identical C2B domains, mainly driven by the polybasic regions 321-332 and arginines 398,399 while interacting with PIP_2_ lipids (see panel 1b). We show that mutations T328E,T329E in the polybasic region disrupt this mechanism of cooperation, drastically increasing the free energy needed to form the stalk.

## 2 Results and Discussion

To induce the formation of the fusion stalk between two initially flat and planar bilayers surrounded by water molecules, we have followed the methodology originally developed by Hub and collaborators ^22,23^ (Prof. Hub generously shared his GROMACS source code with us through personal communications). Using the MARTINI 3 coarse-grained model, we have prepared ternary lipid bilayers containing 1-palmitoyl-2-oleoyl-glycero-3-phosphocholine (POPC), 1-palmitoyl-2-oleoyl-sn-glycero-3-phospho-L-serine (POPS) and the recently developed model for phosphatidylinositol-4-5-bisphosphate lipids (SAP2). This arrangement follows the experimental membrane composition proposed by Jahn and collaborators ^24^ to trap Syt1 to the plasma membrane in the presence of calcium. Accordingly, lipid concentrations were set to POPC:POPS:PIP_2_ (87.5:10:2.5).

The collective variable designed by Hub and collaborators (ξ) induces a hydrophilic trans-membrane pore in a single lipid bi-layer, using a membrane spanning cylinder that is decomposed into slices along the membrane normal ^22^. They also demonstrated that the same collective variable is capable of fusing bilayers under different hydration levels ^23^. Accordingly, in the present work we have used ξ to fuse two bilayers and to study the effects of the Syt1-C2B domain in the process. To facilitate the repeatability of the results and to increase the versatility at user level (and also for our own convenience) we have ported ξ into PLUMED ^25^ as a collective variable (labeled ξ _*f*_).

The source code implementation for the PLUMED environment (including input file parameters and examples) is freely available at https://github.com/lautarodibartolo/MemFusion. See supplementary information for example input files and technical details on the collective variable set of parameters.

Practically, we have used eq. 1 to fuse membranes and form the stalk. The process starts with a pair of flat and parallel independent bilayers (ξ _*f*_ ~ 0.2). Membrane fusion occurs in the interval 0.2 < ξ _*f*_ < 0.85 where the bilayers connect themselves forming the first stalk at ξ _*f*_ ~ 0.58. The collective variable pulls from tail beads (C4A, C4B and C5A) to fill a cylinder with *N_sf_* =85 slices, of thickness *d_sf_* =0.1nm, of radius *R_cylf_* =1.75nm and with an occupation factor ζ_*f*_ =0.5.

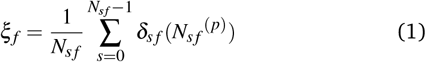

In eq. 1, *N_sf_* ^(*p*)^ accounts for the number of C4A, C4B and C5A beads within slice *s* inside the cylinder. δ*s f* is a continuous function in the interval [0 1] (δ_*sf*_ =0 for no beads in slice *s* and δ_*sf*_ =1 for 1 or more beads in slice *s*). For mathematical details see the original article ^22^.

### 2.1 Hysteresis-free sampling of the fusion stalk

Here, we have equilibrated the inter-membrane distance between bilayers with the necessary amount of cytosolic water molecules to fit one and two C2B domains. Therefore, the PO4:PO4 inter-membrane distance was set to ~3.9 nm, which asks for ~10×10^3^ cytosolic water molecules, imposing a high hydration regime (~34 cytosolic water beads per *nm*^2^). As studied before ^23,26^, different amounts of water molecules between the bilayers result in different equilibrium inter-membrane distances, with significant effects on the free energy landscape for membrane fusion.

To avoid any sampling problems due to the high amount of water molecules we have included in the cytosolic space, we have verified the stalk fusion formation to be hysteresis-free. The free energy cost to evolve from different thermodynamics states (*i.e*. from parallel to fused bilayers) must be independent of the direction of the collective variable ^27^. Therefore, the forward and backward paths from parallel bilayers to the fusion stalk must be identical in the free energy profile. Any differences between them would suggest hysteresis problems, inadequate sampling and poor convergence ^28^.

Accordingly, figure 2a shows PMF calculations in both directions of the collective variable: (i) forwards (black line) and (ii) backwards (red line). Initial configurations in both cases were taken from a slow-growth path in each direction, as originally suggested by Pearlman and Kollman ^29^. These profiles show no significant hysteresis.

**Fig. 2.**
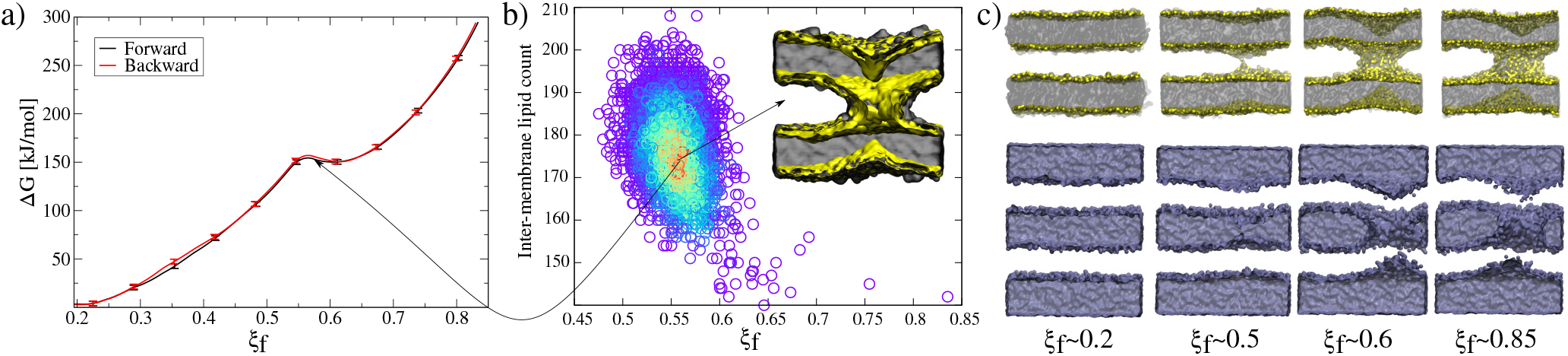
Membranes-only fusion stalk formation. **a)** PMF for membranes-only, forward and backward directions. **b)** Unbiased molecular dynamics showing the local minimum at ξ _*f*_ ~ 0.55. **c)** Molecular dynamics snapshots showing the formation of the stalk. Lipid molecules (top) are shown in grey with PO4 beads in yellow while water molecules (bottom) are blue surfaces.

Along membrane fusion, the first stalk is formed at ξ _*f*_ ~ 0.58 with an energy cost of ~150 *kJ*/*mol*. From that state, the collective variable requires another ~150 *kJ*/*mol* to reach the final state at ξ _*f*_ ~ 0.85 (with a total cost of ~300 *kJ*/*mol*, see figure 2a). Importantly, ξ _*f*_ revealed an energy barrier for the fusion stalk (ξ _*f*_ ~ 0.55) and a local minimum for a metastable stalk (ξ _*f*_ ~ 0.6) (see figure 2a, in agreement with Minimum Free Energy Path (MFEP) dynamics by Smirnova et al. ^30^.

Besides, *μ*s-length unbiased molecular dynamics starting from a well-defined stalk (ξ _*f*_ ~ 0.85) verified the existence of the free energy local minimum at 0.5<ξ _*f*_ <0.6 and though the metastable stalk, see figure 2b. Panel 2c shows molecular dynamics snapshots of the fusion stalk at different stages for lipids only and waters only, separately.

### 2.2 One C2B domain has negligible effects on the fusion stalk free energy profile

Here, using ξ _*f*_ under the MARTINI 3 force-field, with two POPC:POPS:PIP_2_ (87.5:10:2.5) lipid bilayers with ~34 cytosolic water beads per *nm*^2^, we have shown that the necessary work to induce a fusion stalk between initially planar and parallel bilayers is ~300 *kJ*/*mol* (see figure 2a), in good agreement with previous results of similar inter-membrane distances ^5^. Smirnova et al. demonstrated that increasing hydration levels, though larger inter-membrane distances, significantly increase the free energy cost for fusion stalk formation ^26^. Accordingly, Poojari et al. showed for MARTINI POPC bilayers that the energy cost for the fusion stalk is ~175 *kJ*/*mol* for 18 water molecules per *nm*^2 23^.

As figure 3 shows, the introduction of one Syt1-C2B domain in the cytosolic space has little effects on the energy profile, in agreement with our previous study ^5^, (black line for membranes-only and violet line for membranes with one Syt1-C2B). The zero energy reference slightly displaces to the right (from ξ _*f*_ ~ 0.2 to ξ _*f*_ ~ 0.25) as a result of the inward protein pulling from the bilayers. The fusion stalk energy barrier at ξ _*f*_ ~ 0.55 and the local minimum for the metastable stalk at ξ _*f*_ ~ 0.6 are almost identical (between black and violet lines). The total cost for a fusion stalk at ξ _*f*_ ~ 0.85 is slightly lower (~275 *kJ*/*mol*) due to the reduction of the inter-membrane distance (by ~0.2 nm, as measured from unbiased simulations). Importantly, this result is only noticeable by keeping constant the full set of parameters of the collective variable, as adjusted initially for the membranes-only system. As pointed out previously, using different sets of parameters defines different collective variables with different numerical predictions for the free energy ^22^.

**Fig. 3.**
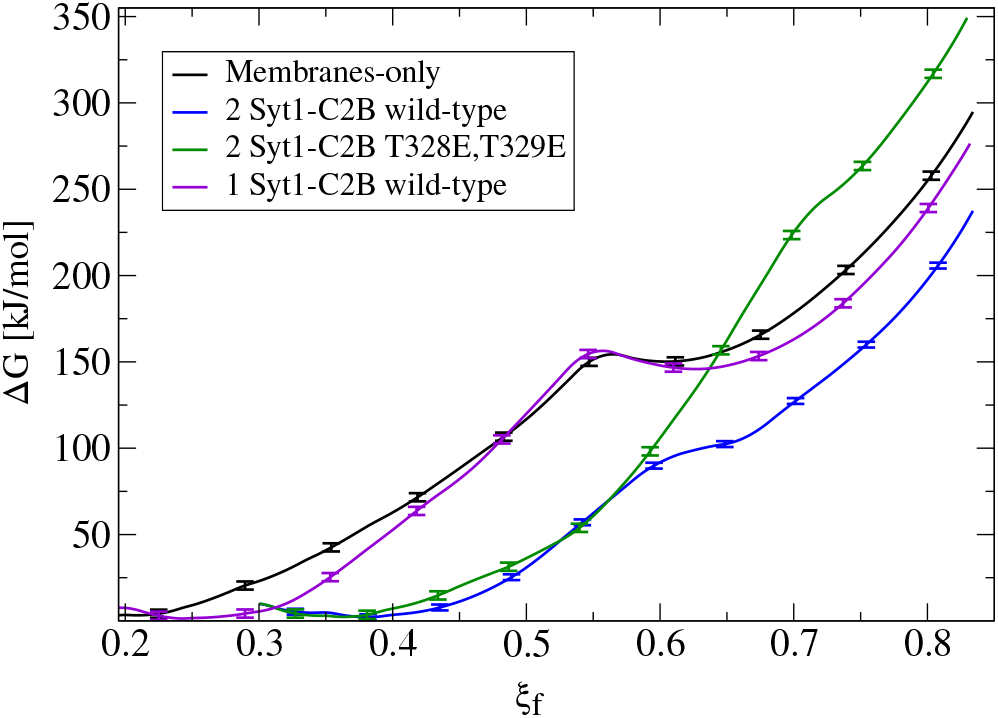
Free energy profiles for the fusion stalk. Membranes-only system (black line) is taken as reference. Repetitions under identical conditions for the same bilayers now containing: one Syt1-C2B wild-type domain (violet line), two wild-type C2B domains (blue line) and two mutant T328E,T329E C2B domains (green line).

### 2.3 Two wild-type C2B domains reduce the free energy cost for the fusion stalk while two mutant C2B domains significantly increase it

Importantly, the presence of two Syt1-C2B domains in the cytosolic space drastically changes the free energy profile. Figure 3 also shows the free energy curve to induce a fusion stalk with two wild-type C2B domains (blue line) and with two mutant T328E,T329E C2B domains (green curve), independently. Scanning mutagenesis, residues 328 and 329 were highlighted by Chapman and collaborators, more than a decade ago, as important for Ca^2+^ triggered fusion ^21^.

For two wild-type domains (figure 3, blue line), it can be observed that both the stalk barrier at ξ _*f*_ ~ 0.55 and the local minimum for the metastable stalk at ξ _*f*_ ~ 0.6 have vanished. Besides, the zero reference is significantly displaced to the right (from ξ _*f*_ ~ 0.2 to ξ _*f*_ ~ 0.35) which lowers the total cost for a fusion stalk at ξ _*f*_ ~ 0.85 to ΔG~240 *kJ*/*mol*. In agreement with this observation, Wu et al. ^31^ recently suggested that Syt1 triggers the opening of an initial fusion pore by bringing the two membranes together and facilitating the exposure of their hydrophobic cores to induce lipid exchange. To verify the displacement to the right of the zero-energy reference, we have performed μs-length unbiased molecular dynamics with two Syt1-C2B domains in the cytosolic space between two planar and parallel bilayers. Under these conditions, the measured equilibrium PO4:PO4 inter-membrane distance was ~1.8 nm, a reduction of 2.1 nm from the membranes-only system (see figure 5b, blue line).

For two mutant domains (figure 3, green line) the curve is similar to two wild-type C2B domains (blue line) until the fusion stalk forms (ξ _*f*_ ~ 0.58). An equivalent displacement to the right of the zero-energy reference is observed, also without the fusion stalk energy barrier or the metastable stalk. Remarkably, the total cost for a fusion stalk at ξ _*f*_ ~ 0.85 has increased to ΔG~350 *kJ*/*mol*, mainly due to the work needed to connect the bilayers. Accordingly, Cafiso and collaborators describe experimentally how mutations lying in the polybasic path of C2B alter the fusion probability ^13^.

### 2.4 Stalk ordering during membrane fusion: from a circular to a square toroid

For the membranes-only system, after the first fusion stalk forms at ξ _*f*_ ~ 0.58, the amount of lipids in the inter-membrane has relatively small variations in the interval ξ _*f*_ ~ [0.58 0.85], see figure 4a (gray circles and black line). This result indicates that the stalk does not widens significantly, although, the evolution from the first stalk to ξ _*f*_ ~ 0.85 is energetically demanding in all cases (see figure 3).

**Fig. 4.**
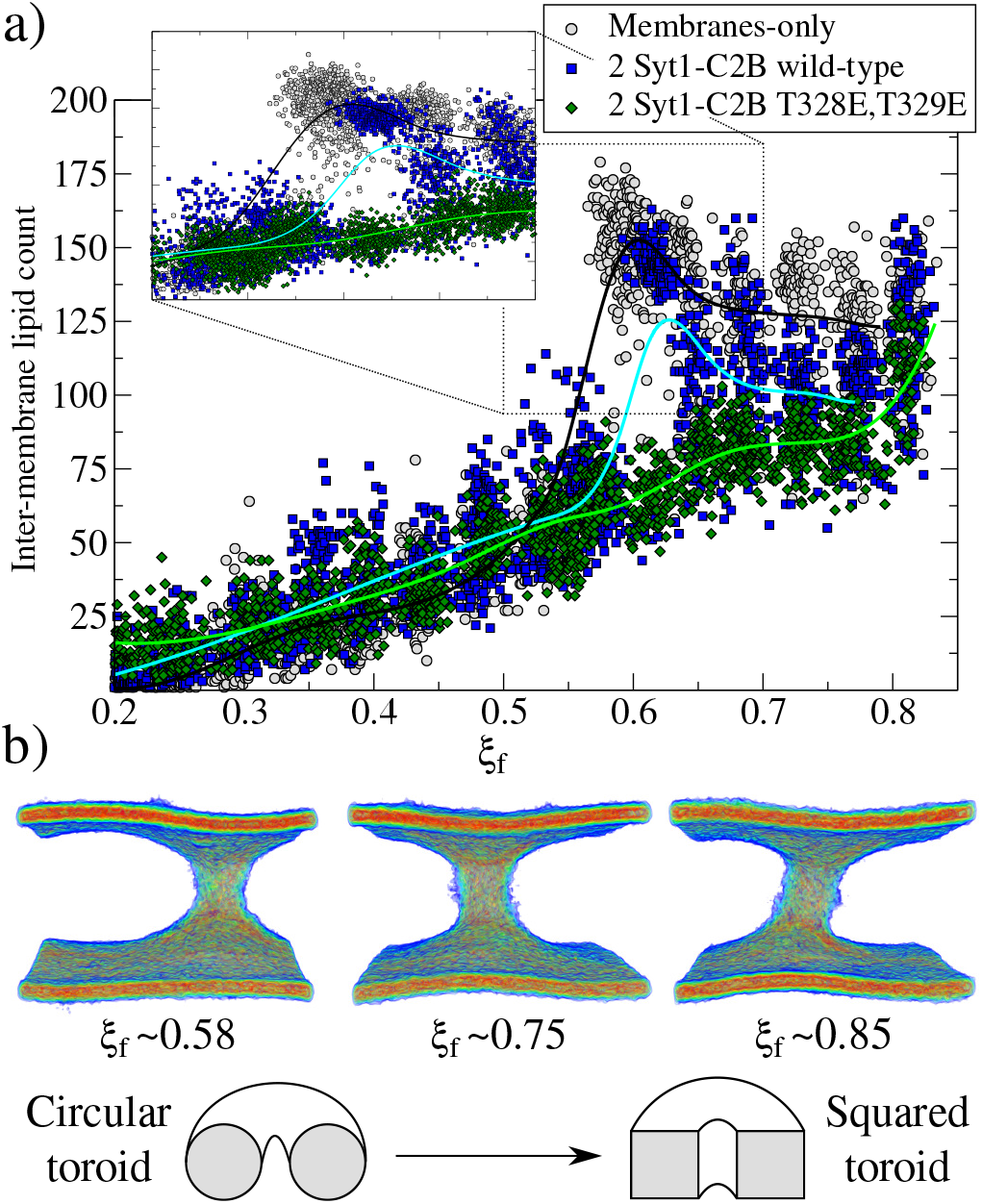
Fusion stalk evolution. a) Inter-membrane lipid count along the evolution of the collective variable ξ _*f*_ for three cases: membranes-only (gray circles), with 2 Syt1-C2B wild-type domains (blue squares) and 2 Syt1-C2B mutant domains (green diamonds). Also, piece-wise interpolating curves are superimposed for each group of data. The inset details the region where the first stalk forms. b) Heat-map colored densities for C4A, C4B and C5A tail beads only, showing the configurational transformation of the toroid for the membranes-only system, as shown by the scheme below.

The reason for almost doubling the free energy with apparently no evident effects is the following: at ξ _*f*_ ~ 0.58 the collective variable has the majority of the necessary lipids to form the stalk already in the inter-membrane space, but they are disordered and not all of them contribute to the collective variable by filling the cylinder slices. As ξ _*f*_ increases (until ξ _*f*_ ~ 0.85) more C4A, C4B and C5A beads from the lipid molecules already in the stalk, order themselves to fill the cylinder slices. Time-averaged densities for C4A, C4B and C5A tail beads only (fig. 4b) show how the geometry of the stalk changes from a circular-toroid at ξ _*f*_ ~0.58 to a square-toroid at ξ _*f*_ ~0.85.

Importantly, systems with membranes-only and containing 2 Syt1-C2B wild-type domains clearly describe the transition at the moment when the first stalk forms (ξ _*f*_ ~ 0.58 for membranes-only and ξ _*f*_ ~ 0.6 for 2 Syt1-C2B wild-type domains). This event is characterized by an accelerated increase in the number of inter-membrane lipids, in agreement with the position of the free energy barrier to form the fusion stalk described in figure 3. Remarkably, for the system containing 2 Syt1-C2B mutant domains the first stalk forms at ξ _*f*_ ~ 0.7 following an almost linear dependency with the amount of inter-membrane lipids, at least right before its final state at ξ _*f*_ ~ 0.85. In the following two sections, we propose a master-servant mechanism of cooperation between C2B domains that explains these behaviors.

### 2.5 The master-servant mechanism (I): polybasic regions 321-332 and arginines 398,399 function as molecular anchors for PIP2 lipids

The mechanism used by wild-type C2B domains to reduce the total work needed to induce a fusion stalk is revealed by specific aminoacid-lipid interactions. The polybasic region in Syt1-C2B domains (321-332) has extensively been studied and is thought to be responsible for crucial interactions with anionic PIP2 lipids ^31–33^, modulating the expansion rate ^12^ and stabilizing the fusion pore ^5^ through PIP_2_ micro-domains at the fusion sites ^5,34^. Moreover, PIP2 clusters have been reported to function as molecular beacons during vesicle recruitment ^32,34^. Recently, Cafiso and collaborators ^13^ have shown that not only the polybasic patch in C2B domains is crucial for membrane fusion but also that arginines 398,399 are key during fusion pore expansion. In their study they demonstrate how the C2B domain makes simultaneous membrane contact with arginines 398,399, the polybasic region and the Ca^2+^ binding loops.

Accordingly, we have performed μs-length unbiased molecular dynamics of 2 wild-type and 2 mutant C2B domains between initially flat and parallel bilayers and we have analyzed C2B spatial orientations. Figure 5a shows histograms for the end-to-end distances along Z axis of each C2B domain. Therefore, higher values of this distance indicate a C2B domain aligned with the normal axis to the bilayers, while lower values suggest a horizontally oriented domain (parallel to the bilayers).

**Fig. 5.**
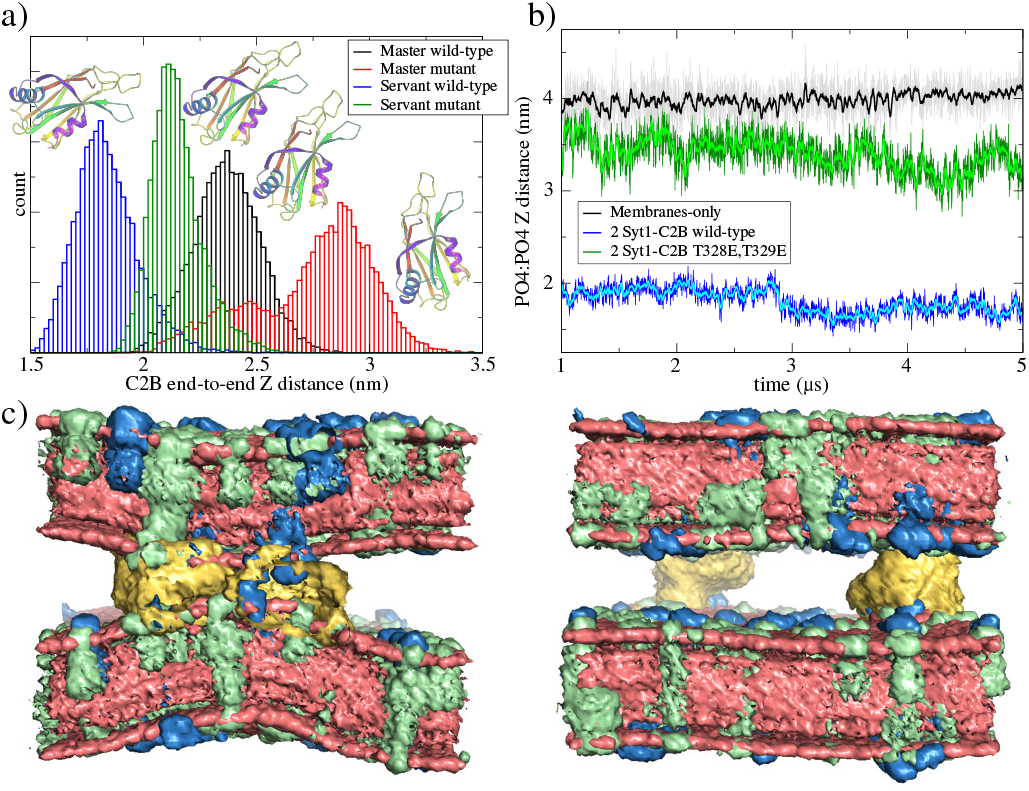
Master-servant C2B domains orientation during unbiased μs-length molecular dynamics. a) Distribution of end-to-end Z distances in C2B domains for two independent simulations performed in a fusion stalk configuration (ξ _*f*_ ~0.85), the first one with 2 wild-type C2B domains (black and blue histograms) and the second one with 2 mutant T328E,T329E C2B domains (red and green histograms). Ribbons representation of the C2B domain schematize the change of orientation observed. b) PO4:PO4 inter-membrane distance for three systems: membranes only (black line), bilayers with 2 wild-type C2B domains (blue line) and bilayers with 2 mutant T328E,T329E C2B domains (green line). c) Averaged densities with POPC in red, POPS in green and PIP_2_ in blue. C2B domains are yellow. Water molecules are not shown. Data collected from μs-length unbiased molecular dynamics started from planar and parallel bilayers.

It can be observed that in average for both independent simulations, one domain (from now on, the master) orients parallel to the Z axis while the other (the servant) is perpendicular to it. For the same unbiased trajectories, figure 5b shows the PO4:PO4 minimum inter-membrane distance for the three systems: membranes-only system (black line), 2 wild-type C2B domains (blue line) and 2 mutant C2B domains (green line). It can be observed that, in the long run, 2 wild-type C2B domains significantly pull membranes together, reducing their PO4:PO4 inter-membrane distance. Importantly, this effect is not observable for 2 mutant C2B domains. Figure 5c shows averaged densities showing the induced curvature by 2 wild-type C2B domains in contrast to almost planar bilayers with 2 mutant C2B domains. Noticeably, 2 Syt1-C2B wild-type domains locally bend the bilayers in around the C2B location. This effect is not observable for 2 mutant domains nor for membranes-only systems. See supplementary figure S6 for a plot measuring minimum inter-membrane PO4:PO4 Z distance as a function of the radial XY distance to the center of the defect, for all three systems under study.

Additionally, we have measured the interactions between the polybasic region (321-332) with PIP_2_ lipids and of arginines 398,399 with the same PIP_2_ lipids, now along two independent μs-length unbiased molecular dynamics one of them containing 2 wild-type C2B domains and the other 2 mutant C2B domains around a fusion stalk of ξ _*f*_ ~0.85. Figure 6 shows radial distribution functions (RDF) for all PIP_2_ lipids in the bilayers alternatively measured from the polybasic region 321-332 (panels 6a and 6b) and arginines 398,399 (panels 6c and 6d). In all cases black lines represent the wild-type C2B domain that directly interacts with the stalk (the master), while blue lines represent the wild-type C2B domain that indirectly interacts with the stalk (the servant).

**Fig. 6.**
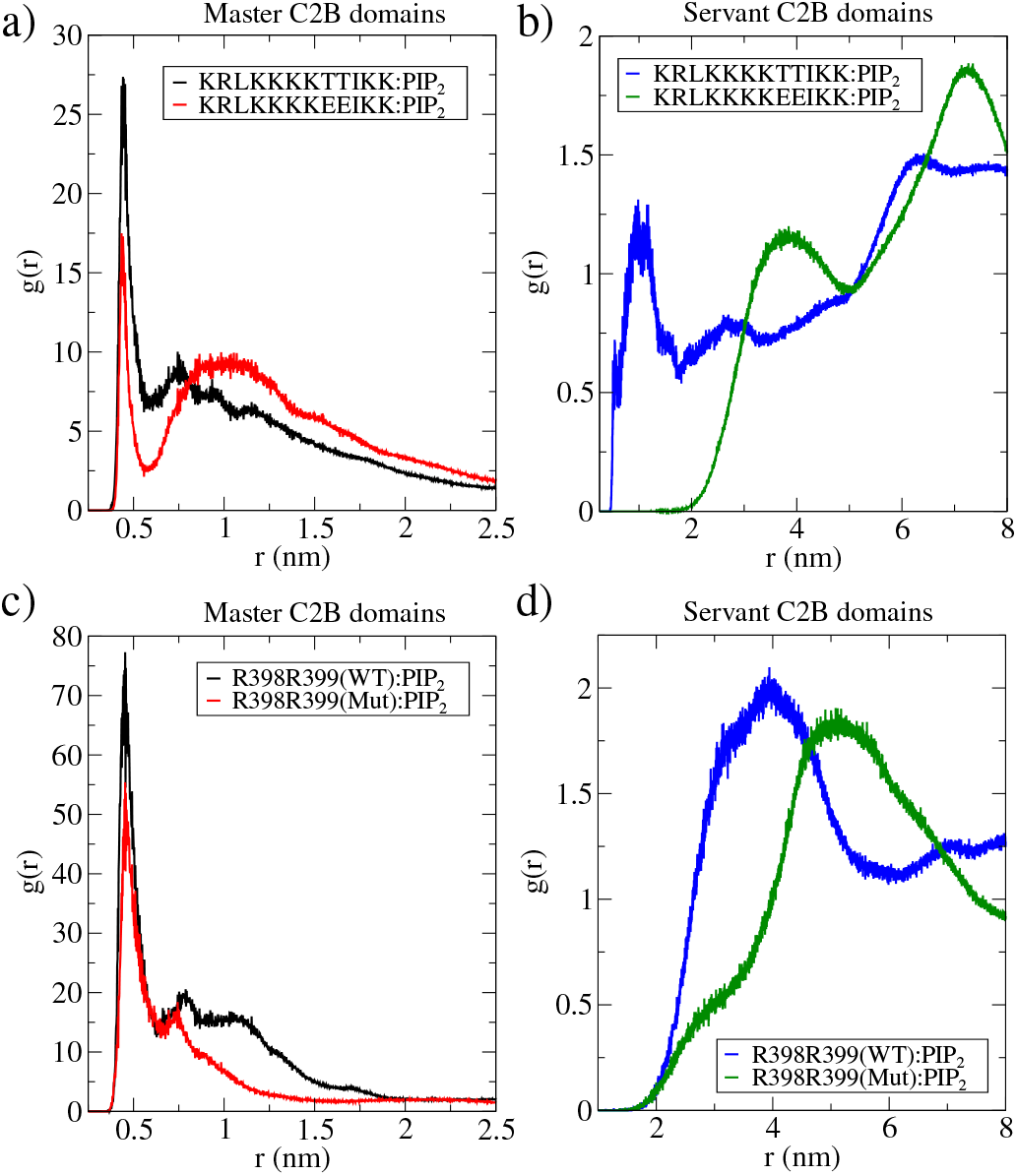
The master-servant C2B mechanism. Radial distribution functions for PIP_2_ lipids measured from the polybasic region (positions 321-332) and arginines (positions 398,399). In all panels, 2 wild-types C2B domains are black and blue lines, and 2 T328E,T329E mutant domains are red and green lines. a) RDF for PIP_2_ with polybasic regions in master proteins (1 wild-type and 1 mutant) as reference. b) RDF for PIP_2_ with polybasic regions in servant proteins (1 wild-type and 1 mutant) as reference. c) RDF for PIP_2_ with arginines R398,R399 in master proteins (1 wild-type and 1 mutant) as reference. d) RDF for PIP_2_ with arginines R398,R399 in servant proteins (1 wild-type and 1 mutant) as reference. Data collected from μs-length unbiased molecular dynamics started at ξ _*f*_ ~ 0.85.

Black lines in panels 6a and 6c show that the master domain highly coordinates with PIP_2_ lipids (peaks at r~0.5 nm) through both its polybasic patch and its arginines 398,399. Simultaneously, blue lines in panels 6b and 6d show that the servant wild-type domain also coordinates well with PIP_2_ lipids through its polybasic patch (peak at r~1 nm) with a less frequent interaction of its arginines 398,399 with PIP_2_ (peak at r~4 nm).

### 2.6 The master-servant mechanism (II): T328E,T329E mutations in C2B domains disrupt the cooperation

The same analysis was applied to 2 mutant C2B domains around an equivalent fusion stalk of ξ _*f*_ ~0.85. Even with T328E,T329E mutations, the mutant master domain (red lines in panels 6a and 6c) shows good interactions with PIP_2_ lipids through both its poly-basic patch and its arginines 398,399, although less frequent than the wild-type master domain (black lines). These results indicate that the main interactions between PIP_2_ and master C2B domains (either wild-type or mutant) is the polybasic region 321-332, in line with previous studies describing PIP_2_ mediated membrane bending ^32,35–37^ and fusion ^36,38–40^. Additionally, also arginines 398,399 appear to be key during C2B:PIP_2_ interactions for both wild-type and mutant master domains, also in agreement with previous data ^13^.

However, servant domains exhibit a different behavior: while the wild-type servant keeps high interactions with PIP_2_ lipids through its polybasic patch, the mutant servant is unable to keep up (see panel 6b). Also, a marginal reduction of arginines 398,399 with PIP_2_ interactions is observed between wild-type and mutant servants (see panel 6d).

## 3 Conclusions

Altogether, these data verifies that C2B has two important regions that interact with anionic lipids, namely the well-known polybasic region KRLKKKKTTIKK (positions 321-332) and the recently described arginines 398,399^13^. We observed a unique behavior of cooperation between identical C2B domains that facilitates the formation of the fusion stalk, as demonstrated by the free energy profile in figure 3 (blue line). While one domain (the master) binds to PIP_2_ lipids though its polybasic region and its arginines 398,399, the other domain (the servant) copies this behavior (see figures 6b and 6d). Together both C2B domains anchor PIP_2_ lipids from different regions to cooperatively reduce the free energy for the fusion stalk to form.

*In-silico* mutagenesis (T328E,T329E) in both C2B domains not only terminates any cooperation to induce the fusion stalk but also increases the associated total work required, making the fusion event thermodynamically more difficult, with respect to the membranes-only system (see figure 3, black and green lines). Remarkably, in the presence of 2 Syt1-C2B mutant domains the formation of the stalk takes place gradually, with a linear dependence on the lipid population of the stalk (see figure 4a, green line). This behavior contrasts with the drastic increase of the amount of lipids in the stalk when 2 Syt1-C2B wild-type domains control the process (blue line).

In terms of its interactions with PIP_2_ lipids, the mutant master domain suffers minor changes with respect to its wild-type counterpart (see figure 6a and 6c), while is the servant mutant domain who is unable to sustain PIP_2_ interactions with neither its polybasic patch nor its arginines 398,399 (see figure 6b and 6d). We propose this reduced interactions as the reason for the whole disruption of the master-servant cooperation mechanism, ultimately responsible for the energetics of the fusion stalk.

## 4 Computational Methods

We have conducted all our simulations using Gromacs-2020.5^41–43^, PLUMED-2.7.2^25^ and the Martini 3 coarse-grained model ^44^. Molecular dynamics simulations used the semi-isotropic NPT ensemble and a time step of 20fs in all cases. The temperature was set to T=303.15K ^6,45–48^ and was controlled by a V-rescale thermostat ^49^ with a coupling constant of 1ps. The pressure was set at 1.0bar with the compressibility equal to 3×10^−4^bar^−1^, using the Parrinello-Rahman barostat ^50^ with a 12ps time constant. Neighbor search used the Verlet cut-off scheme with a buffer tolerance of 0.005kJ/mol/ps and an update-frequency for the neighbor list equal to 25 steps. Periodic Boundary Conditions (PBC) were used in all directions. Coulomb interactions used the reaction field method with a cut-off of 1.1nm and a relative dielectric constant of 2.5. Van der Waals interactions followed the cut-off scheme set to 1.1nm.

In all cases, we have used a pair of lipid bilayers containing 1024 molecules each. These bilayer patches of ~17×17nm ensure negligible finite-size effects due to interactions between periodic images of the fusion pore ^5,51^. In all cases the pair of bilayers were solvated in more than ~30×10^3^ W coarse-grained water molecules to fulfill the ample water condition for MARTINI ^52^. The PO4:PO4 inter-membrane separation was adjusted to equilibrate at ~3.9nm to fit one and two Syt1-C2B domains. This inter-membrane distance results in ~10×10^3^ W water beads in the cytosolic space. PIP_2_ lipids for MARTINI 3 were modeled following the parametrization by Melo and collaborators ^53^ (https://github.com/MeloLab/PhosphoinositideParameters).

Figures were created using Visual Molecular Dynamics (VMD) ^54^, the academic version of Maestro ^55^, Grace (GRaphing, Advanced Computation and Exploration of data) ^56^, Inkscape ^57^, GIMP (GNU Image Manipulation Program) ^58^ and Gnuplot ^59^. Averaged densities from molecular dynamics simulations were generated using GROmaρs ^60^ and PyMOL ^55^.

### 4.1 PLUMED implementation of the collective variable

We have implemented the collective variable to induce membrane fusion as a modular C++ file compilable with PLUMED. The file is freely available at https://github.com/lautarodibartolo/MemFusion together with a README file and an example input system. Additionally, in supplementary information we have included an example input file for PLUMED-2.7.2 to induce the fusion stalk using the same POPC:POPS:PIP_2_ bilayers described in this work.

### 4.2 PMF calculations

Free energy profiles were computed with umbrella sampling ^61,62^ in PLUMED ^25^ and recovered with using the Weighted Histogram Analysis Method (WHAM) using the implementation developed by Prof. Grossfield ^63^. Fusion stalk free energy profiles required 15 windows to span ξ _*f*_ in the interval [0.2,0.85] using a force constant k=30,000 kJ/mol. All windows used to recover the free energy profile contained at least 100ns in the steady-state regime, although the total simulation time required for each window varied (from 110ns to 210ns) depending on the region of the profile. See supplementary information for convergence analysis on the free energy profiles and for technical details on umbrella sampling windows distribution.

### 4.3 MARTINI coarse-grained

PMF calculations with the new MARTINI 3 are in good agreement with previous results using the MARTINI 2.2 version ^5^. A slight difference in the length of the pre-fusion region (ξ _*f*_ <0.58) was although observed. Under MARTINI 2.2, using the previous model of PIP_2_ lipids (POP2) and a different set of parameters, the first fusion stalk forms later in the collective variable space (0.65<ξ _*f*_ <0.7). This effect could also suggest that MARTINI 3 is prone to membrane fusion and though less thermodynamic work would in principle be needed to fuse bilayers, as also pointed out by Hub and collaborators ^23^. Importantly, Vanni and collaborators have recently shown that MARTINI 3 is particularly suitable for characterizing mutagenesis experiments in peripheral proteins while binding to lipid bilayers ^64^.

## Author Contributions

DM conceived the idea and supervised the research. DM and ALDB designed the computational experiments and performed the numerical simulations. ALDB wrote the C++ PLUMED implementation of the code in consultation with DM. DM wrote the manuscript in consultation with ALDB. Both authors discussed the results and the conclusions.

## Acknowledgements

Supercomputing time for this work was provided by CCAD (Centro de Computación de Alto Desempeño de la Universidad Nacional de Córdoba). Grants from CONICET (PIP-13CO01) and ANPCyT (PICT2017-1002) are gratefully acknowledged as well as GPU hardware granted by the NVIDIA Corporation. The authors thank Prof. Jochen Hub for generously sharing his GROMACS source code with us, Prof. Giovanni Bussi for his useful advice in PLUMED C++ coding and Dr. Mariano Polo for critically reading the final manuscript.

