## Supplementary Information for "Synaptotagmin-1 C2B domains cooperatively stabilize the fusion stalk via a master-servant mechanism"

### Umbrella sampling technical details

We performed umbrella sampling using PLUMED. All windows contain at least 100ns in their steady-state. Transient regimes were individually discarded and additional simulation time (between 10ns and 100ns) was added. Equilibrium points ($\xi_{f0}$) for each window were distributed as follows: 0.20, 0.25, 0.30, 0.35, 0.40, 0.45, 0.50, 0.55, 0.58, 0.60, 0.65, 0.70, 0.75, 0.80, 0.85. In all cases a force constant of k=30,000 kJ/mol was used. Figure S1 shows convergence of the energy profile in the membranes-only system. As observed, there are no significant differences between 90ns and 100ns. Besides, figure S2 shows histograms of the order parameters for all windows. Figure S3 shows time-averaged densities during stalk formation for selected values of the collective variable.

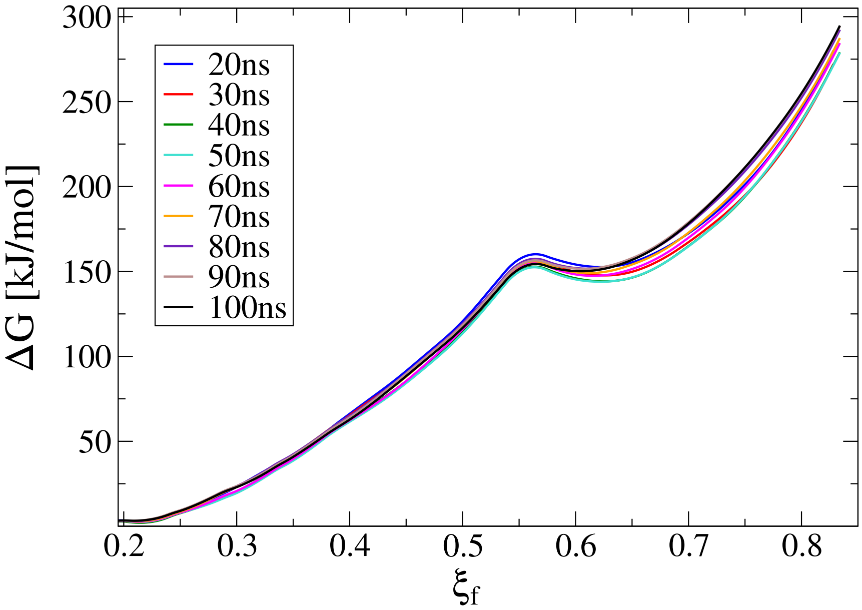


Figure S1: **Umbrella sampling convergence analysis.** Free energy profile for membranes-only at different simulation times in the steady-state regime.

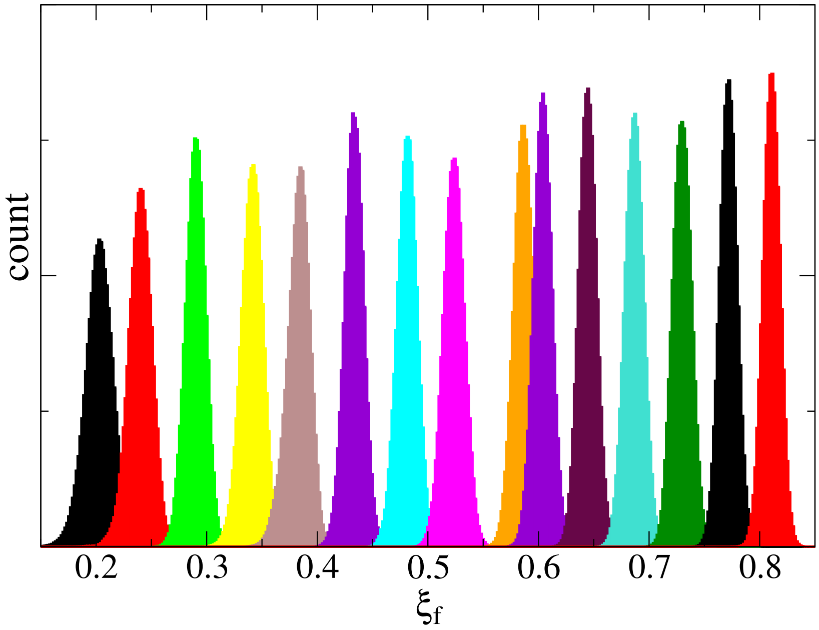


Figure S2: **Histogram superposition for umbrella sampling simulations**. A set of 15 windows was used to recover free energy profiles in figure S1.

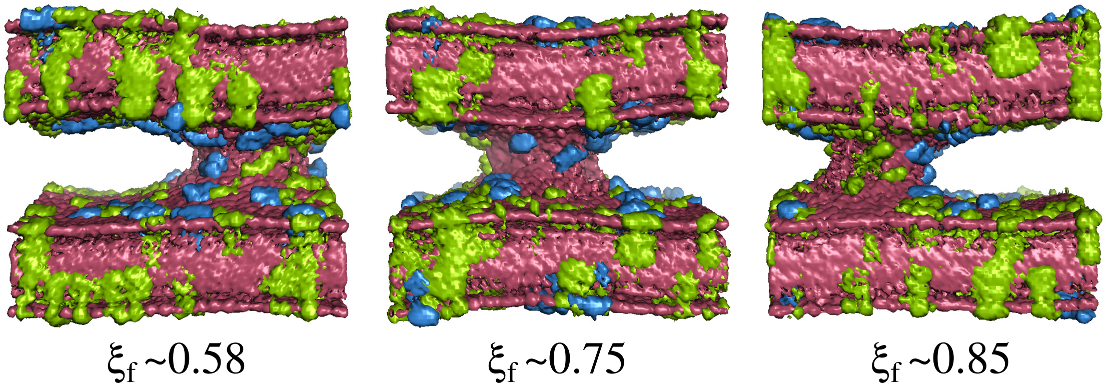


Figure S3: **Time averaged densities during stalk evolution**. Colors are assigned as follows: POPC in red, POPS in green and $PIP_2$ in blue. Water molecules are not shown.

### Unbiased molecular dynamics: from initially flat and parallel bilayers

We performed 5$\mu$s of unbiased molecular dynamics starting from initially flat and parallel bilayers, under the three cases studied in this work: membranes-only, with 2 Syt1-C2B wild-type domains and with 2 Syt1-C2B mutant domains. Figure S4 shows molecular dynamics snapshots at t=1$\mu$s, wild-type domains are blue and mutant ones are green.

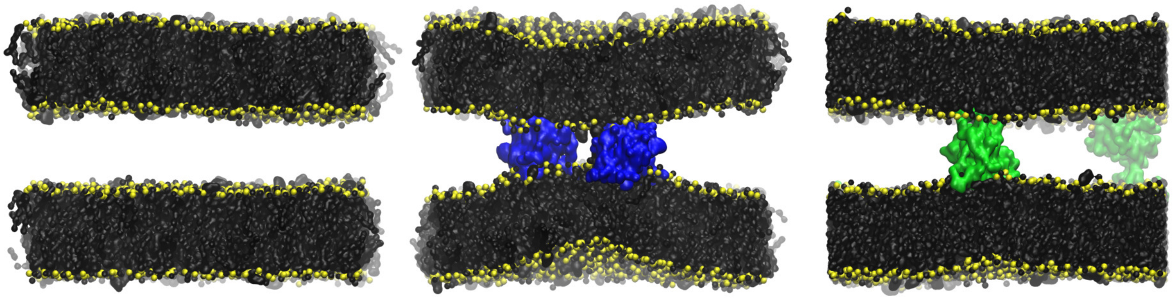

Figure S4: **Molecular dynamics snapshots at t=1$\mu$s.** Lipid molecules are shown in black with PO4 beads in yellow. Wild-type Syt1-C2B domains are blue and mutant ones are green.

Figure S5 shows lipid density profiles for the systems containing 2 C2B domains (wild-type and mutant). From all three panels, $PIP_2$ lipids shows the more significant difference between wild-type and mutant systems. This observation verifies that wild-type C2B domains mainly pull from $PIP_2$ lipids to bring membranes together. In agreement with Radial Distribution Functions (RDF) in figure 6 in the main text.

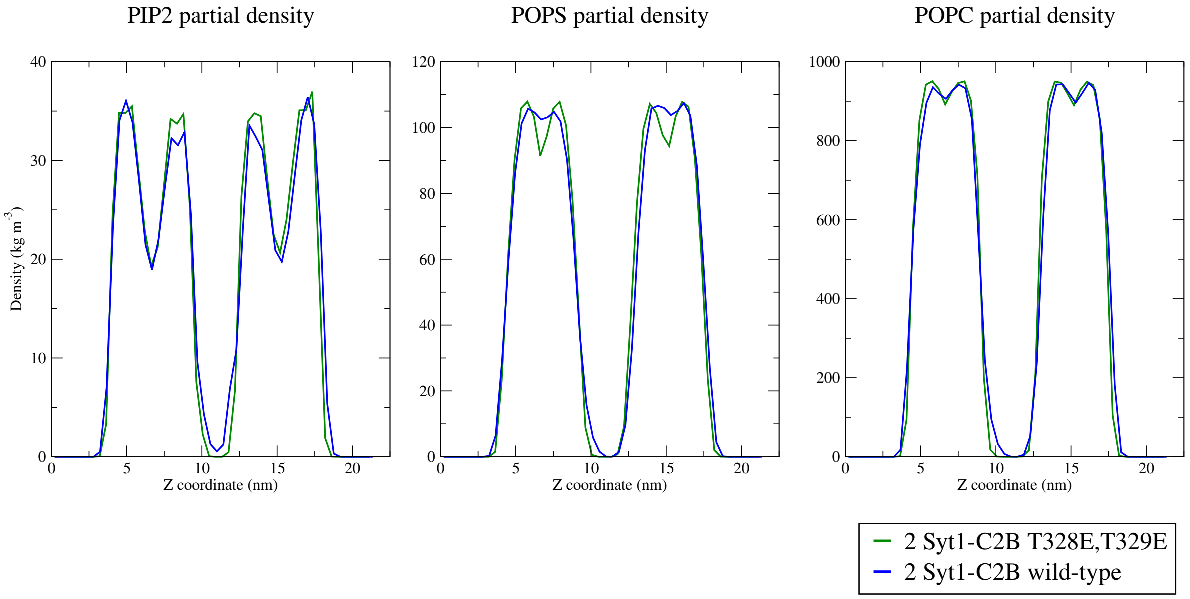


Figure S5: **Lipid species density profiles.** Profiles show systems with 2 Syt1-C2B wild-type domains (blue) and with 2 Syt1-C2B mutant domains (green).

Figure S6 shows the minimum PO4:PO4 distance along Z axis as a function of the radial XY distance to the center of the defect (where most probably the stalk will form). It can be observed that 2 Syt1-C2B wild-type domains (blue line) locally pulls membrane together to a minimum inter-membrane distance of ∼1.5nm. On the contrary, both membranes-only and mutant domains systems keep the inter-membrane separation above 3nm, as already suggested by molecular dynamics snapshots in figure S4.

Figure S6 also verifies that the defect is initially a local deformation, meaning that all curves converge to ∼4.25nm for radial distances far enough from the defect (∼6.5nm).

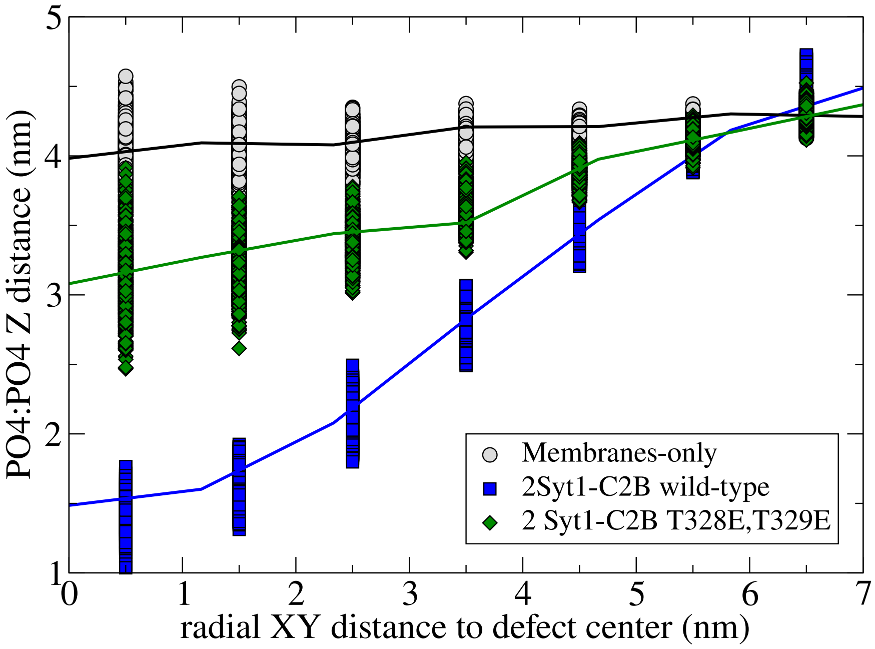


Figure S6: **PO4:PO4 Z distance as a function of the radial XY distance to the center of the defect.** Data shows membranes-only system (black line), with 2 Syt1-C2B wild-type domains (blue) and with 2 Syt1-C2B mutant domains (green).

### Unbiased molecular dynamics: from a $\xi_f \sim 0.85$ fusion stalk

We performed 10$\mu$s of unbiased molecular dynamics starting from a fusion stalk initially set at $\xi_f \sim 0.85$. Figure S7 shows the inter-membrane amount of lipids. Both curves describe how the fusion stalk stabilizes at ∼175 lipids. This amount of lipids is in agreement with figure 2b in the main text. Also, when the stalk is released from its restraint ($\xi_f \sim 0.85$ at t=0$\mu$s in figure S7) under both situations (with 2 wild-type and 2 mutant domains) the amount of lipids in the inter-membrane space increases. Accordingly, initial values in this figure are in agreement with the amount of lipids measured under the collective variable restraint in figure 4a in the main text.

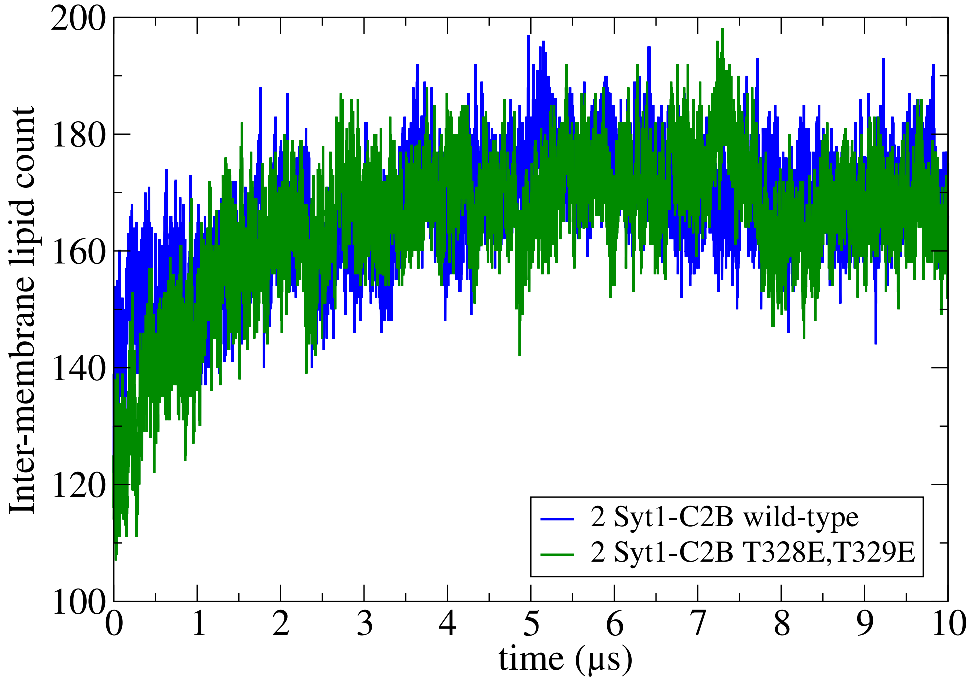


Figure S7: **Inter-membrane lipid count.** Measured for systems containing 2 wild-type Syt1-C2B (blue line) and 2 mutant Syt1-C2B domains (green line).

### PLUMED: input file to induce a stalk at $\xi_f \sim 0.85$

```
INCLUDE FILE=groups.dat

Xi1: FUSIONPORE1 UMEMBRANE=uMem LMEMBRANE=lMem TAILS=tails NSMEM=85 DSMEM=0.1 ..

.. HMEM=0.25 RCYLMEM=1.75 ZETAMEM=0.5

restraint: RESTRAINT ARG=Xi1 AT=0.85 KAPPA=30000.0

PRINT ARG=Xi1 FILE=COLVAR085 STRIDE=1
```

### PLUMED: groups.dat

```
lMem: GROUP ATOMS=1-10752,21505-22728,23953-24420

uMem: GROUP ATOMS=10753-21504,22729-23952,24421-24888

tails: GROUP ATOMS=8-23948:12,12-23952:12,23966-24884:18,23970-24888:18
```
